# Biofabrication using maize protein: 3D printing using zein formulations

**DOI:** 10.1101/2020.07.29.227744

**Authors:** Jorge Alfonso Tavares-Negrete, Alberto Emanuel Aceves-Colin, Delia Cristal Rivera-Flores, Gladys Guadalupe Díaz-Armas, Anne-Sophie Mertgen, Plinio Alejando Trinidad-Calderón, Jorge Miguel Olmos-Cordero, Elda Graciela Gómez-López, Esther Pérez-Carrillo, Zamantha Judith Escobedo-Avellaneda, Ali Tamayol, Mario Moisés Alvarez, Grissel Trujillo-de Santiago

## Abstract

The use of three-dimensional (3D) printing for biomedical applications has expanded exponentially in recent years. However, the current portfolio of 3D printable inks is still limited. For instance, only a few protein matrices have been explored as printing/bioprinting materials. Here, we introduce the use of zein, the primary constitutive protein in maize seeds, as a 3D-printable material. Zein-based inks were prepared by dissolving commercial zein powder in ethanol with or without polyethylene glycol (PEG400) as a plasticizer. The rheological characteristics of our materials, studied during 21 days of aging/maturation, showed an increase in the apparent viscosity as a function of time in all formulations. The addition of PEG 400 decreased the apparent viscosity. Inks with and without PEG400 and at different maturation times were tested for printability in a BioX bioprinter. We optimized the 3D printing parameters for each ink formulation in terms of extrusion pressure and linear printing velocity. Higher fidelity structures were obtained with inks that had maturation times of 10 to 14 days. We present different proof-of-concept experiments to demonstrate the versatility of the engineered zein inks for diverse biomedical applications. These include printing of complex and/or free-standing 3D structures, materials for controlled drug release, and scaffolds for cell culture.

## Introduction

The biofabrication community is actively seeking to harness the advantages of 3D printing for solving biomedical challenges; however, several factors are still preventing the full exploitation of this new printing technology. The advantages of traditional extrusion-based 3D-printing include layer-by-layer fabrication of almost any 3D form at relatively high resolution, down to the microscale [1]. For instance, 3D printing has enabled the fabrication of free-form products, such as personalized dosed pills, temporary stents, tissue-engineered heart valves or full-sized bladders [2–6]. However, the diversity of biomedical devices that can be 3D printed is severely limited by the portfolio of available printable materials or inks. Printing will be further enabled by expanding the spectrum of extrudable biofriendly materials (i.e., biocompatible, resorbable, amenable to biological functionalization, etc.) [7,8]. Among the many possibilities, protein-based materials are particularly attractive. Proteins are constitutive blocks in the human body; therefore, their use may lead to the design of biocompatible, resorbable, cell-friendly, and bio-smart inks for biomedical applications.

Today, only a limited number of protein matrices have been explored as printing/bioprinting materials. Relevant examples are mammal-derived proteins, such as collagen [9], fibrin [10] or gelatin [11]. Some of these materials (i.e., collagen and gelatin) are extensively used in bioprinting applications because of their cell adhesive and bioactive properties that allow cell attachment or interaction. Their mechanical properties are also similar to those of natural tissues, making them suitable host environments for cultured cells [11–13]. However, these proteins have limited availability and typically lack the physical stability needed for 3D printing of precise and large structures (cm range) [8]. Furthermore, their use often requires tight control over the processing temperature, chemical pre-functionalization such as methacrylation, and post-printing cross-linking steps [14–16]. These processability challenges are further complicated by the limited availability of mammal-derived materials and, thus, their high cost [11,17].

Fortunately, alternative 3D-printing materials can be found outside the mammal group. For instance, silks from arthropods and insects exhibit excellent characteristics of biocompatibility and stability under physiological conditions and has been confirmed as 3D printable, thereby greatly enhancing its potential in biofabrication [18]. In the present study, we explore the use of zein, the most abundant constitutive protein in maize seeds, as a suitable material for developing 3D printing materials. Zein, as a byproduct of corn syrup or cornstarch production, is a widely abundant and cost-effective plant-derived biopolymer [19,20]. It is a prolamine, a protein rich in the amino acid proline, and occurs as aggregates linked by disulfide bonds [21,22]. It has a molecular weight around 25,000 to 35,000 kDa and is poorly soluble in water due to its relatively high content of hydrophobic amino acids (proline, leucine, and alanine); however, it also exhibits amphiphilic behavior due to its high glutamine content (21–26%) [19,23].

Zein is biodegradable, edible, and has been classified as “generally recognized as safe” (GRASS) by the FDA [24]. It has been used in biomedical applications, such as the coating for pharmaceutical capsules [19,20,23] or the fabrication of scaffolds for cell culture [25]. To our knowledge, zein has never been explored as a 3D printing material, even though concentrated solutions exhibit shear thinning properties [26], which are highly favorable for extrusion-based 3D printing. Here, we explore the use of concentrated zein solutions in ethanol/water as inks for 3D printing.

Zein-based formulations, optionally added with polyethylene glycol (PEG-400) as a plasticizer [25,27], were characterized in terms of their rheology, printability, and stability. We also report the effect of ink aging time on the rheological properties and printability. The shape fidelity and self-standing behavior of zein-based inks was successfully demonstrated by the 3D-printing of complex multilayer constructs. Furthermore, we show proof-of-concept experiments that demonstrate some potential applications of our formulations. For instance, we 3D printed zein (Z) and zein-PEG (ZP) antibiotic-loaded tablets and studied their release kinetics and bacterial inhibition properties. We also seeded 3D printed zein scaffolds with C2C12 myoblasts to assess cytocompatibility.

## Materials and methods

### Materials

Zein (Z3625, SLBV3020), poly-(ethylene glycol) 400 (P3265, MKBG3641V), Dulbecco’s Modified Eagle’s Medium (DMEM; D5748, SLBW4140), Dulbecco’s phosphate buffered saline (PBS; D5773, SLBW277), nalidixic acid (NA; N8878, BCBW6556), and 4′,6-diamidino-2-phenylindole (DAPI; D9542, 28718-90-3) were purchased from Sigma Aldrich, USA. Mouse myoblast cells (C2C12) and *Staphylococcus aureus* (ATCC^®^ 29213) were purchased from ATCC, USA. Fetal bovine serum (16000-044, 2087367) and antibiotic-antimycotic 100X (15240062) were purchased from Gibco (Grand Island, USA). Phalloidin-iFluor 647 (ab176759, GR3256003-6) was purchased from Abcam. PrestoBlue^®^ (A13261, 2044809) was purchased from Invitrogen. Luria-Bertani (LB) broth medium was purchase from Difco, MD, USA.

### Zein ink preparation

We prepared two different zein ink formulations. A pristine zein ink (Z) was prepared by solubilizing 15 g of zein in 20 mL of 70% ethanol aqueous solution. This mixture was homogenized by vortexing for 10 to 15 minutes, then heated at 45 °C for 20 min under manual and gentile agitation and stored (when still hot) in tightly closed sterile 10-mL syringes to avoid further solvent evaporation and water uptake. The zein-PEG ink (ZP) was prepared using the described protocol but incorporating 2 g of PEG400 into the mix of 15g zein in 20 mL of 70% of ethanol aqueous solution. The PEG400 was added during the first ten minutes of heating. The rheological and printability characterization of these inks was conducted after 1, 7, 10, 14, and 21 days of storage/aging time.

### Rheological characterization of zein inks

The rheology of the zein inks was evaluated using a MCR 500 rheometer (Anton-Paar Rheoplus, Austria) equipped with a Peltier device for temperature control. A 50 mm parallel plate geometry and a gap of 5 mm were used for all measurements. The effective viscosity, the storage modulus (G’) and the loss modulus (G”) were measured using an angular frequency of 10 rad s-1 and a shear rate between 0 and 0.5 s^−1^ at room temperature at different aging/maturation times (*i.e.*, 4, 7, 10, 14, and 21 days). Three independent repeats were performed (n=3).

### Printability studies

We evaluated the printability of zein-based formulations by printing a grid design at different combinations of relevant printing variables, including extrusion pressure, printing velocity, and ink maturation time. Three settings of pressure (65, 75, and 85 kPa), printing velocity (3, 5, and 7 m s^−1^), and ink maturation times (4, 7, 10, and 14 days) were explored in our printing experiments. All 3D-printing experiments were conducted using a BioX (Cellink, Sweden) bioprinter. Cartridges (3 mL volume) were filled with Z and ZP inks aged for 4, 7, 10, or 14 days. Blue high-precision conical 22 G nozzles (410 μm internal diameter) from Cellink were used for printing. Four-layer reticles (grids) of 2 cm × 2 cm were extruded and immediately photographed after printing. At least six evaluations of the ISL value (n=6) were performed per grid.

### Characterization of the fidelity of the printed structures

We evaluated the fidelity of grid structures printed with zein-based inks using image analysis techniques. Photographic images of printed structures were analyzed using ImageJ 1.52q software. The heights and widths of the empty spaces between each zein line were measured by transforming the photographs into 8-bit images and modifying their contrast to facilitate the detection of borders. A set of 18 lines for each group was obtained and averaged. The average inner square length (ISL) values were graphed with their corresponding standard deviations (Figure 3B).

We also assessed the effect of the extrusion temperature (at the nozzle) on the printability of zein-based inks. To do this, we fabricated grids of Z and ZP formulations at temperatures of 10, 25, and 45 °C using a pressure setting of 75kPa and a linear printing velocity of 3mm/s.

### Fourier transform infrared spectroscopy analysis

Fourier-transform infrared spectroscopy (FTIR) analysis was performed to evaluate the effect of aging and water exposure on the protein secondary structures of zein-based inks.

FTIR spectra of 7- and 14-day aged Z and ZP inks, before and after immersion in water, were obtained using a FTIR/FT-NIR Perkin Elmer Spectrum 400, equipped with a total attenuated reflection (ATR) accessory. Sixteen scans per sample at 4 cm^−1^ resolution were acquired over the range of 1100–1900 cm^−1^. A secondary-structure analysis was performed by band-narrowing techniques, second-derivative analysis, and Fourier self-deconvolution using the Origin software (Version 2019). In particular, the spectral region from 1570–1720 cm^−1^, corresponding to the amide I region, was analyzed following the protocol described by Yang et al. [28]. Bands with a frequency of 1650–1658 cm^−1^ were assigned to α-helixes, from 1640–1650 cm^−1^ to non-ordered structures or random coils, from 1660–1680 cm^−1^ to loop or β-turns, and from 1620–1640 and 1670–1695 cm^−1^ to β-sheets [28].

### Constructs stability under aqueous conditions

We also evaluated the stability of 3D-printed zein grids in culture medium. Briefly, we incubated grids in DMEM cell culture medium at 37 °C. The thickness of the lines for the grids printed using Z and ZP inks were measured over a period of 7 days using brightfield microscopy. Thickness was normalized according to the following equation:

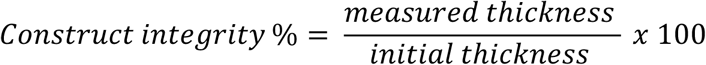

We also printed spiral structures at 50 kPa and 20 mm/s and submerged them in water to explore the behavior of the zein-based printed constructs exposed to different aqueous environment scenarios. Some spirals were printed and submerged in water at 25 °C for 30 minutes immediately after printing. Others were desiccated for 48 h after printing and then submerged in water at 25°C for 30 minutes. Structures were photographed immediately after recovery from the water bath. This experiment was conducted with three technical repeats (n=3).

### Drug release studies

We evaluated the release of an antibiotic from 3D-printed Z and ZP tablets loaded with NA. Briefly, a solution of 45 mg NA in 40 mL 70% ethanol was added during the formulation of zein inks. The Z and ZP inks were prepared as described previously while maintaining temperatures at 37 °C during dissolution to preserve the NA antibiotic activity. Homogenous solutions containing 3 mg NA per g zein were obtained by vortexing. Cylindrical tablets 5.6 mm in height and 10 mm in diameter were printed using the BioX printer and dried in a desiccator for 3 days. The printed tablets of Z and ZP were placed in 15 mL Falcon tubes containing 10 mL PBS and left to float freely at 37 °C and pH=7 in a water bath to study drug-release kinetics under physiological conditions. Throughout the drug release experiments, 1 mL aliquots of PBS were withdrawn for analysis and fresh PBS was added to maintain a constant volume. The release of NA into the medium was determined by measuring absorbance at 258 nm using a Nanodrop 1000 spectrophotometer (Thermo Fisher Scientific, USA). We used a calibration curve to correlate the absorbance with NA concentrations in the range of 0.5 to 500 μg μL^−1^. Three independent experiments (N=3) were conducted with three technical repeats (n=3).

### Antibiograms

We performed assays to evaluate the antimicrobial activity of NA released from 3D-printed zein tablets on *Staphylococcus aureus* strains. The bacteria were cultured in tryptic soy broth overnight at 37 °C under continuous shaking until the optical density (OD) reached 0.6–0.8 (at 600 nm) [29,30]. Tablet dissolution methods were used to determine the antimicrobial effect of 3D-printed discs according to antimicrobial testing guidelines [31]. Briefly, for broth dilution experiments, 1 mL MHA broth was placed in Falcon tubes and 100 μL of bacterial suspension was added. The 3D-printed tablets of different zein formulations (i.e., Z, ZP, Z-nalidixic, and ZP-nalidixic) were then placed into the tubes. Tubes were shaken at 100 rpm and 37 °C for 48 h until the final reading at 600 nm. Tests were performed in triplicate (n=3).

In addition, we conducted convenctional antibiograms in Petri dishes. Briefly, bacterial dilutions containing 10^7^ colony forming units (c.f.u.) of *S. aureus* were plated in solid plate count agar and incubated at 37 °C in the presence of 3D-printed cylindrical zein tablets (of 5.6 mm in height and 10 mm in diameter). Each tablet contained ~1.32 mg NA. The area and diameter of the inhibition halo was measured 72 hours after seeding using image analysis techniques.

### Cell scaffolding

We conducted cell culture experiments using Z and ZP 3D-printed grids as cellular scaffolds. We extruded 2-layer grids (0.81 mm height) in a BioX 3D bioprinter and dried them in a desiccator for 7 days. The grids were then sterilized under UV light overnight and then placed in 6-well ultra-low attachment plates and washed repeatedly with PBS to remove any embedded ethanol. In a first round of cell-culture experiments, grids were surface seeded with C2C12 myoblast cells (ATCC CRL 1772) by covering the grids with a C2C12 cell suspension containing 5 × 10^5^ cells mL^−1^ and incubating for 24 hours at 37 °C to promote attachment. Non-attached cells were removed after this incubation period. We ran parallel cell culture experiments in conventional 6-well plates as a control. Cultures were maintained in 3 mL DMEM culture medium supplemented with 10% FBS and 1% antibiotic-antimycotic, and incubated at 37 °C in a 5% CO_2_ atmosphere for 7 days. The grids were observed at days 0, 1, 3, and 7 using an Axio M2 Observer fluorescence microscope (ZEISS, Germany). PrestoBlue^®^ metabolic activity assays were performed at day 0, 1, 3, and 7. Samples at day 7 were stained with phalloidin (1:1000 in PBS) and DAPI (1:1000 in PBS) at 4°C overnight, washed 3 times with PBS, and observed under the fluorescence microscope. Two independent experiments (N=2) with triplicates (n=3) were conducted.

In a second round of cell culture experiments, 3D-printed Z and ZP grid scaffolds were incubated in 50 mg mL^−1^ fibronectin in PBS overnight at 4°C to enhance cellular attachment. After incubation, the fibronectin solution was removed and the grids were placed in fresh ultra-low adhesion well plates. C2C12 cells were seeded over the grids and cultured for 7 days, as described before. Cell attachment and spreading were observed by fluorescence imaging with phalloidin and DAPI staining on day 7. All experiments were conducted in triplicate (n=3).

### Scanning Electron Microscopy

We analyzed the internal microstructure of the different zein ink formulations using scanning electron microscopy (SEM). A line of zein ink was extruded using a syringe and subsequently stored in a desiccator for 7 days at room temperature. Cross-sections of the dried constructs printed with zein inks aged for 7, 14, and 21 days were exposed by manual breaking. These cross-sections were coated with gold using a Q150R ES rotating sputter machine (Quorum, United Kingdom) prior to imaging. SEM micrographs were obtained at 500× magnification in a ZEISS EVO MA25 SEM (Germany). Three technical repeats (n=3) and three images per sample were analyzed.

The number and projected area of the zein pores were measured using ImageJ from the SEM images. Results were analyzed in MATLAB R2018B to evaluate the evolution of pore size distribution through aging in zein formulations with or without PEG400. The pore size values were grouped in 12 bins spanning the range from 0.2 μm^2^ to 15.32 μm^2^. We reported pore size distributions and cumulative frequency plots based on these calculations.

### Statistical analysis

ANOVA analysis was performed with Minitab Express 1.5 (2017). Differences with a *p* value < 0.05 (*) were considered statistically significant.

## Results and discussion

Here, we introduced the use of zein, the primary constitutive protein in maize, as a 3D-printable material. Two different ink formulations were prepared by dissolving commercial zein powder in ethanol (Z), and optionally adding polyethylene glycol (PEG400) as a plasticizer (ZP). Figure 1A schematically shows the method of preparation of these zein inks. Briefly, zein powder was dispersed in ethanol by vigorous vortexing. The resulting suspension was heated at 45 °C until a homogeneous solution was obtained. For our ZP formulations, PEG400 was added during the first ten minutes of heating. Z and ZP formulations were characterized in terms of rheology and printability.

**Figure 1.**
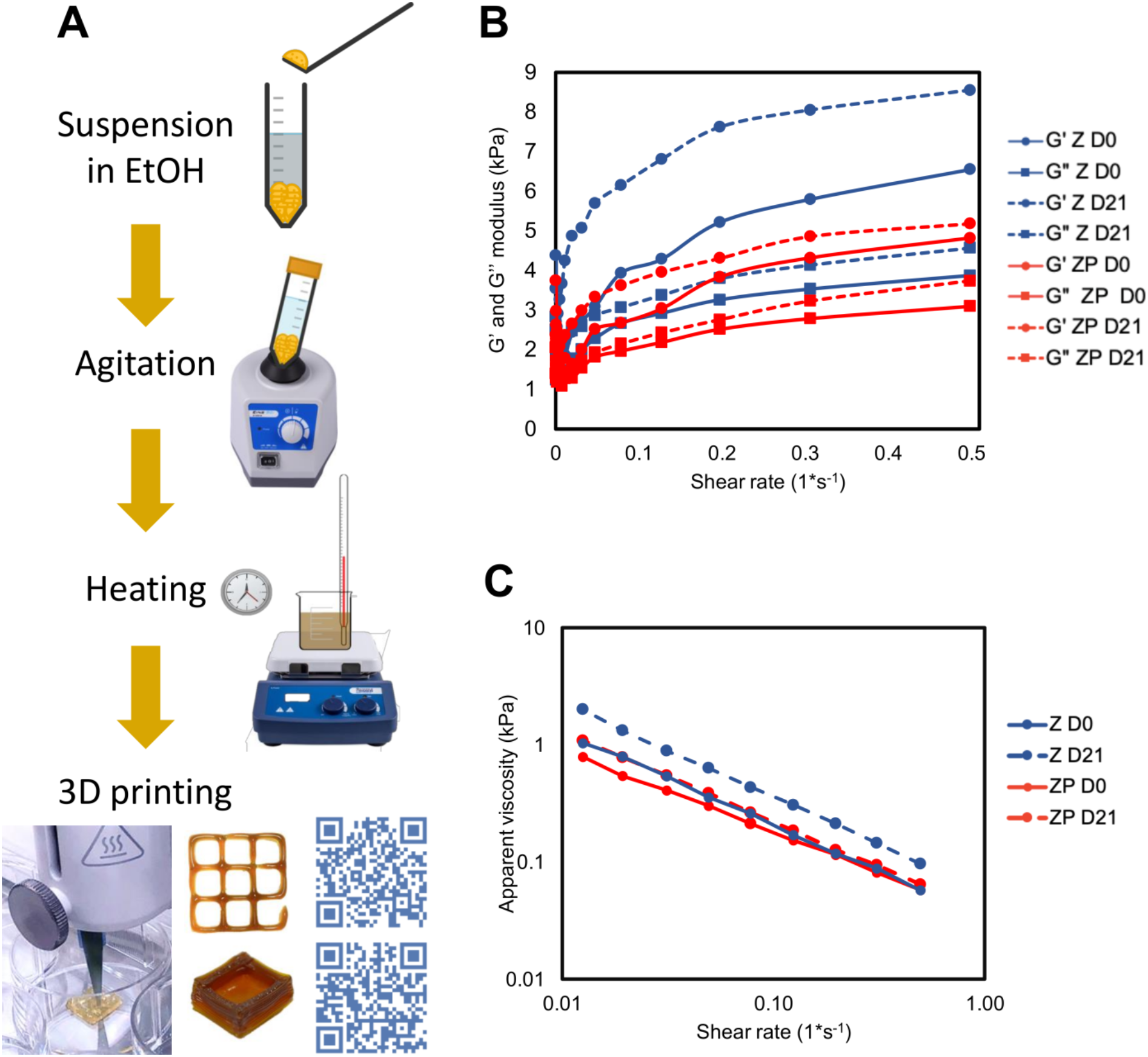
Preparation and characterization of zein inks for 3D printing. A) Schematic representation of the protocol for preparation of zein-based inks. Zein powder was added to an aqueous ethanol solution and gradually incorporated by vortexing to generate a suspension, which was subsequently heated to produce a homogeneous solution. These formulations were placed in tightly closed syringes and stored for 21 days (See Supplementary video 2). (B-C) Rheological characterization of zein (Z) and zein + PEG400 (ZP) inks was conducted at day 0 and 21: B) Storage (G’) and loss modulus (G’’) and C) apparent viscosity of zein formulations at room temperature as a function of shear rate (n=3).

One of the main findings of our work is that important properties of these zein formulations, as related to their printability attributes, evolve with aging time. We characterized the evolution of these properties (i.e., viscosity, viscoelastic behavior, and printability itself).

### Rheology characterization

In extrusion printing, the rheology of an ink greatly determines printability [32–34]. We have conducted a detailed rheological characterization of our zein ink formulations at the time of preparation and after 21 days of aging. The following observations were derived from this characterization.

Both zein formulations (Z and ZP) behave as pseudo plastics with G’>>G’’ (Figure 1B). This pseudoplastic behavior is maintained over the maturation time of 21 days. Consistently, the storage modulus (G’) is higher than the loss modulus (G’’) over the whole frequency range, indicating a typical gel behavior [35]. Frequency oscillation experiments showed that storage (G’) and loss modulus values (G’’) increased as the Z and ZP ink matured over 21 days. For example, the storage modulus for Z formulations increased from 6.55 ± 0.14 kPa at day 0 to 8.55 ± 0.09 kPa at day 21 when evaluated at the highest shear rate. For ZP formulations, the storage modulus at the highest shear rate increased from 4.82 ± 0.12 kPa at day 0 to 5.18 ± 0.07 Pa at day 21.

Our inks behave as shear thinning materials, meaning that their effective viscosity decreases with increasing shear rates. Remarkably, the viscosity of our zein inks decreases by more than one order of magnitude within the window of shear rates that we analyzed (Figure 1C). Overall, shear thinning behavior is a key characteristic sought in inks for extrusion 3D printing [36,37] as it strongly impacts printability. In practical terms, shear thinning inks will exhibit less resistance to flow as more mechanical (or hydraulic) pressure is applied during extrusion. The incorporation of PEG does not alter the shear-thinning behavior of zein inks. As expected, PEG addition yields inks with a lower viscosity and a lower storage modulus [38]. The addition of PEG decreases viscosity, but the slope of the viscosity versus shear rate plot does not change (Figure 1C).

Ink viscosity is highly dependent on maturation time and PEG attenuates this effect. For example, at shear rate values in the neighborhood of 0.01 s-1, the viscosity of Z inks increases by approximately 1 kPa with respect to the value on the day of preparation. However, ZP inks exhibit a much lower increment of viscosity (~ 0.3 kPa) after 21 days of storage.

The aging effect in zein solutions has been attributed to structural changes in the protein [26,39]. For instance, the aging effect has been related to changes in secondary structure over time, as analyzed by small-angle X-ray diffraction [40]. A full mechanistic understanding underlying the changes in viscosity in zein inks through aging is beyond the scope of this work; however, we conducted basic FTIR spectroscopy analysis to identify possible structural variations in zein formulations through aging based on the vibrational amide I bands related to the protein secondary structures [41]. We estimated the changing percentages of different structural configurations (i.e., α-helix, β-sheets, β-turns, and non-ordered structures or random coils) in the overall structure of zein formulations with aging. The percentage of each secondary structure is available as supplementary information (Figure S1 and Table S1). The results suggested that the α-helix content of the Z formulation increased by 2.06% as aging progressed (Supplementary Table S1). By contrast, the proportion of β-sheets and non-ordered structures increased in the aged ZP formulation by 13.34%, and 21.08%, respectively, while the percentage of α-helix decreased by 7.69%. During aging, the Z and ZP formulations show a decrease in the content of loop structures (−5.27% for Z and −26.73% for ZP) and an increase in non-ordered structures (5.22% for Z and 21.08% for ZP). These observations suggest that the change in viscosity may be induced by changes in the protein structure during aging and that the rate and thermodynamics of these changes might be modified by PEG addition [26,40].

### Printability of zein formulations

Our formulations were printable in a wide range of conditions in a commercially available 3D bioprinter. Printability, or the ability of an ink to be extruded in a manner that creates well-defined structures, is commonly evaluated in terms of the morphology, shape fidelity, and self-standing ability of printed constructs [42]. We evaluated the printability of Z and ZP formulations using a grid design printed with inks at different maturation times. In addition, a matrix of conditions that included different extrusion pressures and printing-head linear velocities was explored. Figure 2 shows that aging time played a crucial role in the printability of our inks. Freshly prepared Z and ZP inks (at 4 days of aging) were poorly printable under all tested conditions. Their viscosity was too low to render defined grids; instead, a high amount of extruded material formed squared puddles. Low viscosity, as determined at the early stages of maturation, correlates well with the low fidelity of printings conducted using 4- and 7-day aged inks. The higher viscosities of the 10- and 14-day aged inks resulted in improved fidelity. As aging time and viscosity increased, grids with higher definition were obtained (Figure 2 and 3).

**Figure 2.**
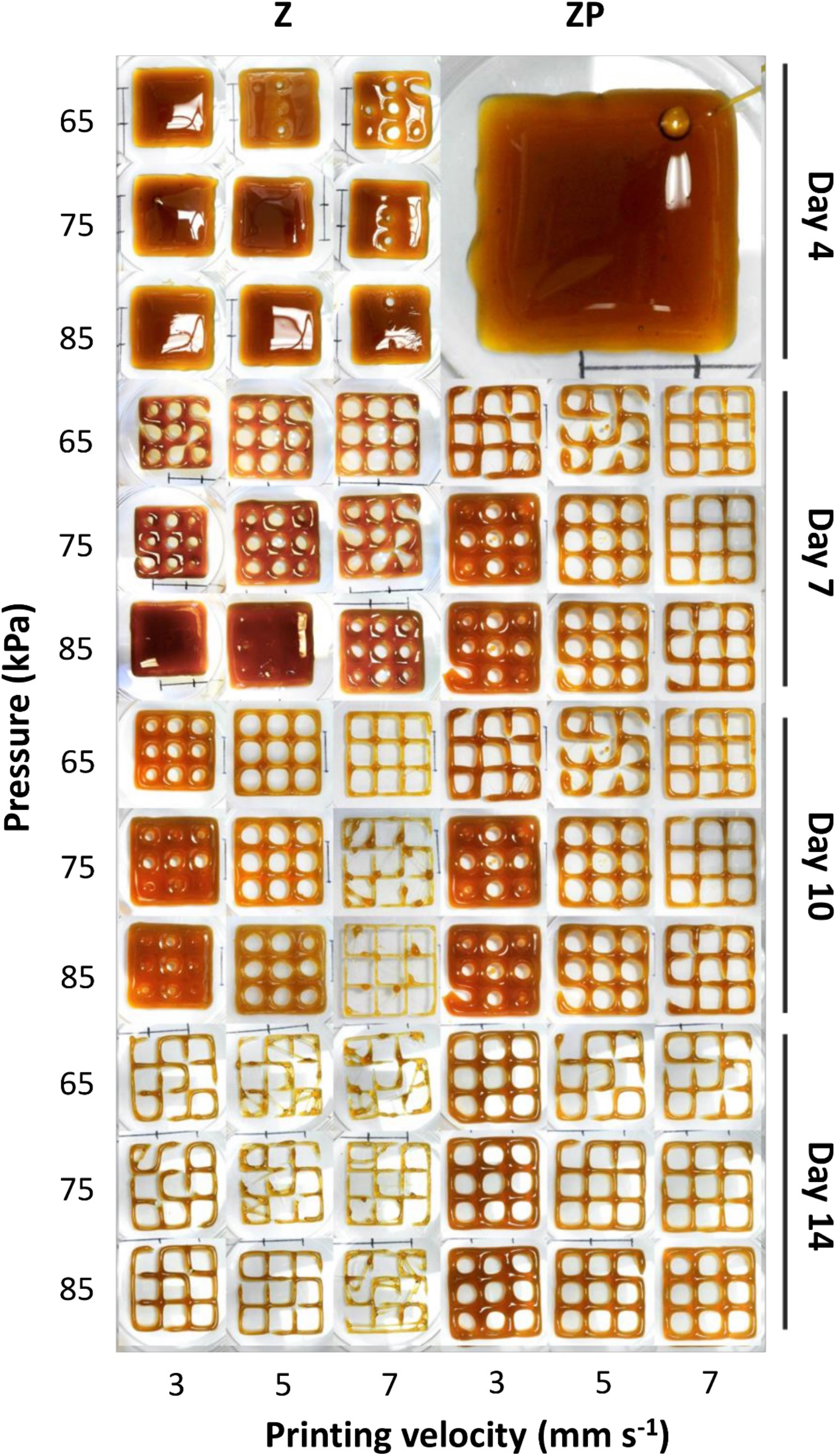
Effect of maturation time on the printability of Z and ZP inks. Different maturation times (i.e., 4, 7, 10, and 14 days), linear printing velocities (i.e., 3, 5, and 7 mm s^−1^), and extrusion pressures (i.e., 65, 75, and 85 kPa) were explored (n=3). Scale bar: 1cm.

**Figure 3.**
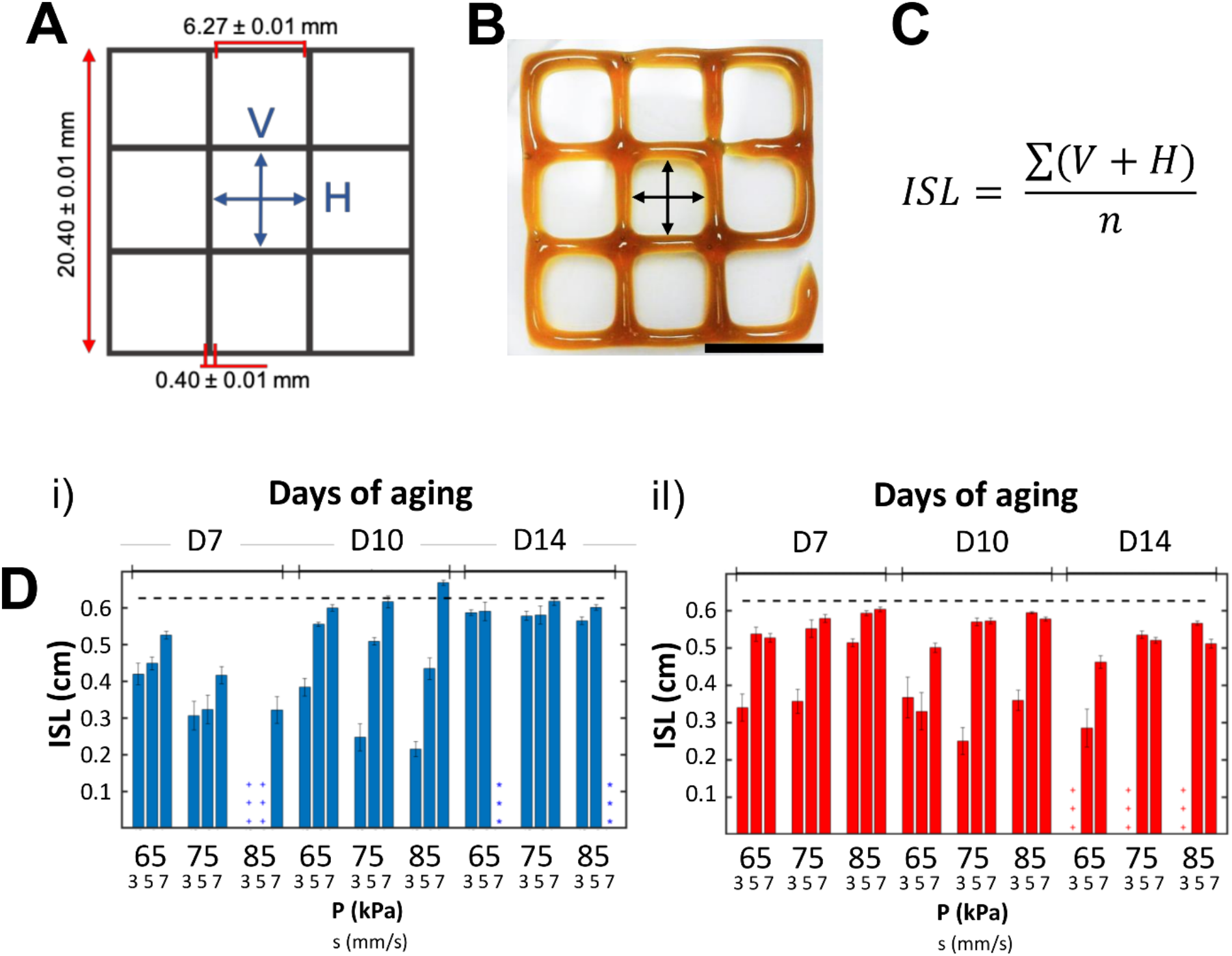
Analysis of the fidelity and quality of our printed zein structures. A) STL grid design. The theoretical dimensions of vertical (V) and horizontal (H) lines is shown (*i.e*., V and H = 0.627 cm.) B) Actual printed grid; vertical and horizontal lines are indicated. Scale bar: 1 cm. C) Equation used to calculate the internal square length (ISL). D) ISL values as determined for different printing conditions. Grids printed with zein (Z) and zein-PEG formulations (ZP) were analyzed. ISL values for (i) Z (blue bars), and (ii) ZP (red bars) grids printed at different pressure and speed conditions. Each triplet of bars shows variations depending on linear speed. Analysis is repeated for inks aged 7 (D7), 10 (D10), and 14 (D14) days. Cases in which printing parameters were inadequate to generate continuous and defined structures are marked with a set of tree asterisks (***). When the linear velocity of the printing-head was too fast or the pressure was higher than needed, a cross (+++) was assigned.

Results also showed a clear interplay between ink rheology and printing conditions (linear speed and pressure). Since matured inks (aged >14 days) were thicker, they required higher pressures and lower velocities to achieve good printability. A clear effect of the plasticizer (PEG400) was also evident in our inks. The incorporation of PEG 400 as plasticizer to zein formulations has been reported before, and an effect was expected [25,43]. Pristine zein inks fail to develop continuous grids at the printing conditions used, while ZP inks performed well and produced structured reticles in practically all conditions tested. Incorporation of PEG400 also extended the printability time frame. ZP developed coherent structures only after 14 days of aging, while Z inks were printable from day 7 at low pressures (65 and 75 kPa) and high velocities (5 and 7 mm s^−1^).

We conducted a simple quantitative analysis of the quality and fidelity of our printed structures. This analysis was based on a simple measurement of the distance between opposite parallel lines in printed grids of zein (Figure 3A-B), and comparing the average of these distances among different printing conditions.

The average of these distances, referred as the inner square length (ISL), is an indicator of fidelity (Figure 3C). ISL analysis offers a more objective view of the influence of printing parameters on printing quality in a real multivariable scenario. Ideally, a single extruded line should be as thick as the nozzle gauge from which it was printed. Therefore, deviations from the designed ISL value are an indicator of deviations from ideal printing quality. For instance, in our experiments, the space between each parallel line in the STL file is 6.27 mm, considering a line thickness of 0.4 mm (diameter of the nozzle outlet) (Figure 3A). In a high-fidelity printing process, the measured ISL should be equal to 6.27 mm, in perfect agreement with the STL design of our grids (Figure 3A).

For a particular printing condition, the standard deviation (STD) of the ISL value relates to reproducibility. Smaller values of STD indicate higher reproducibility. The printing quality among different printing parameters can be evaluated by comparing the measured ISL with the target (i.e., intended ISL) value of 6.27 mm. Figure 3D summarizes the results obtained from this quantitative ISL analysis. Within this set of results, general trends of printing quality, as related to the printing parameters, are readily evident. For instance, Figure 3D clearly shows that, for a particular formulation, fidelity tends to increase with aging. Note that discontinuous structures or cuts in the printed structures due to bioprinting lapses, delays, or interruptions during extrusion were observed under several printing conditions. These printing discontinuities can be attributed to the interplay of rheological and printing parameters. For example, lower linear velocities allow high-viscosity inks to be deposited with precision and avoid material dragging by the printhead. However, when the printing linear speed is lower than ideal, the material forms clumps in some spots and thinner lines in others, thereby increasing the STD of ISL values. In some cases, the ISL of the printed grid is higher than the ideal value. This happens mainly because some layers are not correctly printed due to a printing speed that is faster than the flow of the material.

Our printing quality assessment identified a further set of additional observations. Printability is evidently a strong function of pressure, and an adequate selection of the pressure setting is key for zein 3D printing. However, the pressure range for feasible printing is wider than that of the speed range. In other words, our results suggest that the printing speed for zein can affect the deposition of material in a more sensitive way than is observed for pressure. In general, at constant pressure values, the higher the printing speed (within the range of speeds tested), the better the grid fidelity achieved (Figure 3D i, ii).

In addition, aging increases viscosity, and higher viscosities ease the control required to achieve high printing quality. Note, however, that the effect of aging in printability is more evident in pristine zein inks than in those with added PEG400. In general, 14-day aged Z inks exhibit lower ISL than their 10-day aged counterparts, whereas the printability of ZP inks is statistically similar after 10 or 14 days of maturation. As expected, the fluidity is also higher for ZP than for Z ink formulations. This observation is consistent with the lower viscosity conferred by the addition of PEG400 to the ink formulations. The ZP inks are only printable after 14 days of aging, whereas some Z formulations exhibit good printability after 11 days of aging.

### Influence of printing temperature on zein ink printability

Temperature strongly influences ink viscosity; therefore, the extrusion temperature setting is another relevant parameter that determines printability. Here, we evaluated the printability of 14-day aged Z and ZP inks at different extrusion temperatures of 10, 25, and 45°C (see Figure 4) [25]. The Z and ZP inks responded differently to temperature. Printability of high-viscosity inks (i.e., inks with no PEG400 or aged inks) is favored at high printing temperatures, while the printability of low-viscosity inks (i.e., those containing PEG400 or unaged inks) favors lower temperatures. Extrusion of Z inks at low temperatures reveals that their increase in viscosity hinders their ability to flow and, therefore, to be successfully printed. However, the printability of younger or PEG400-containing (less viscous) inks is improved when using low temperatures, even at 10 °C [44]. By contrast, high temperatures are useful for printing aged inks.

**Figure 4.**
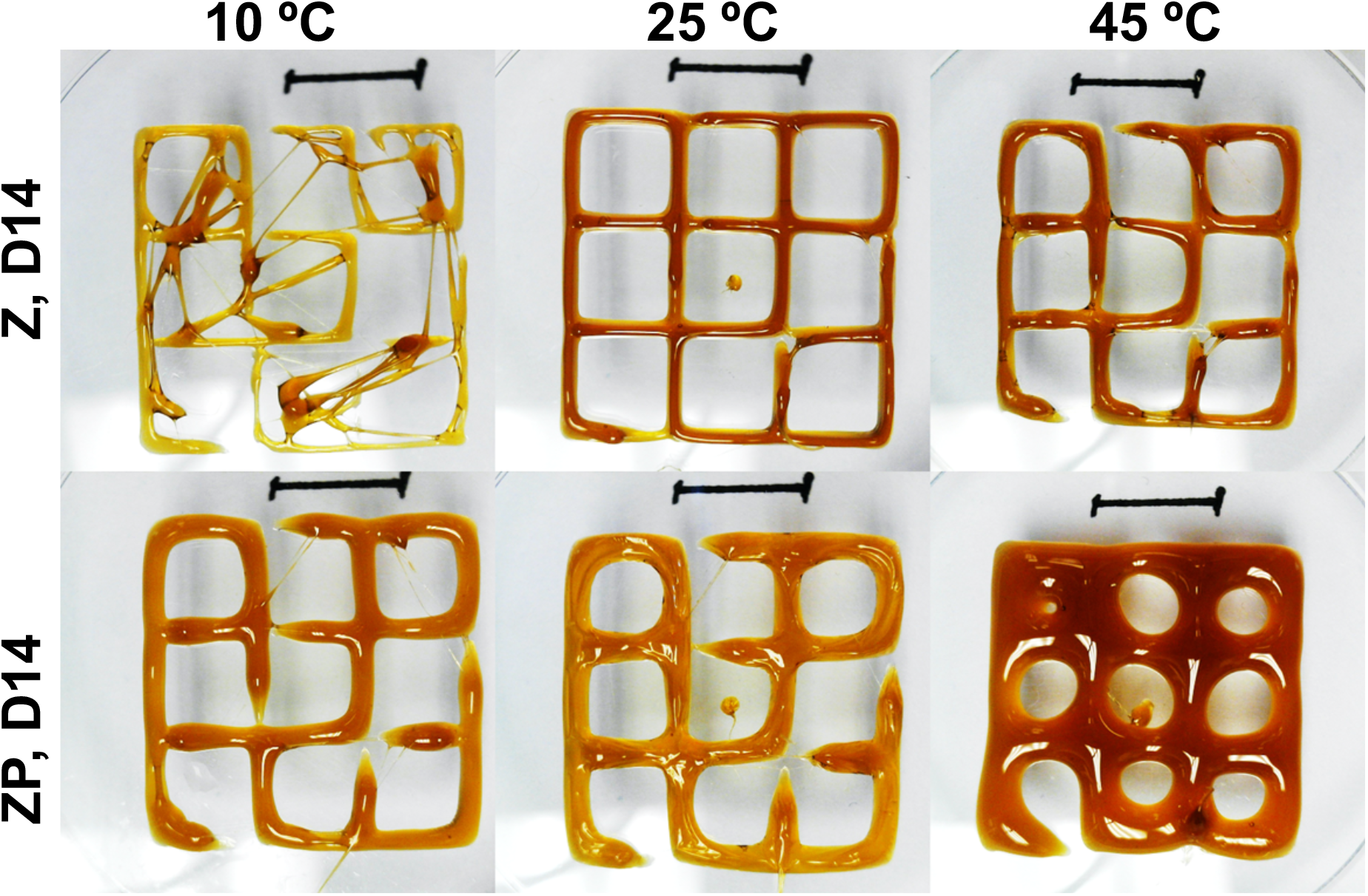
Temperature effect on zein ink printability. Z and ZP inks 14-day old printed at 10, 25, and 45°C at 75kPa and 3mm/s

Overall, our results suggest that zein formulations may be tuned to accommodate different printing aims. By knowing the range of capacities of a particular bioprinter, one can choose an adequate formulation for a particular profile of pressures and speeds. Even for a fixed composition, the rheology of a zein-based ink can be fine-tuned by playing with different maturation times, PEG concentrations, and printing temperatures.

### 3D printing of complex structures

We also demonstrated the capability of zein-based inks to produce complex 3D structures. Figure 5A shows the Tecnológico de Monterrey logo and Figure 5B shows a close-up of the fine features printed using zein-based inks. The logo had a length of 13.3 cm and a height of 1.6 mm and was built using four layers of material.

**Figure 5.**
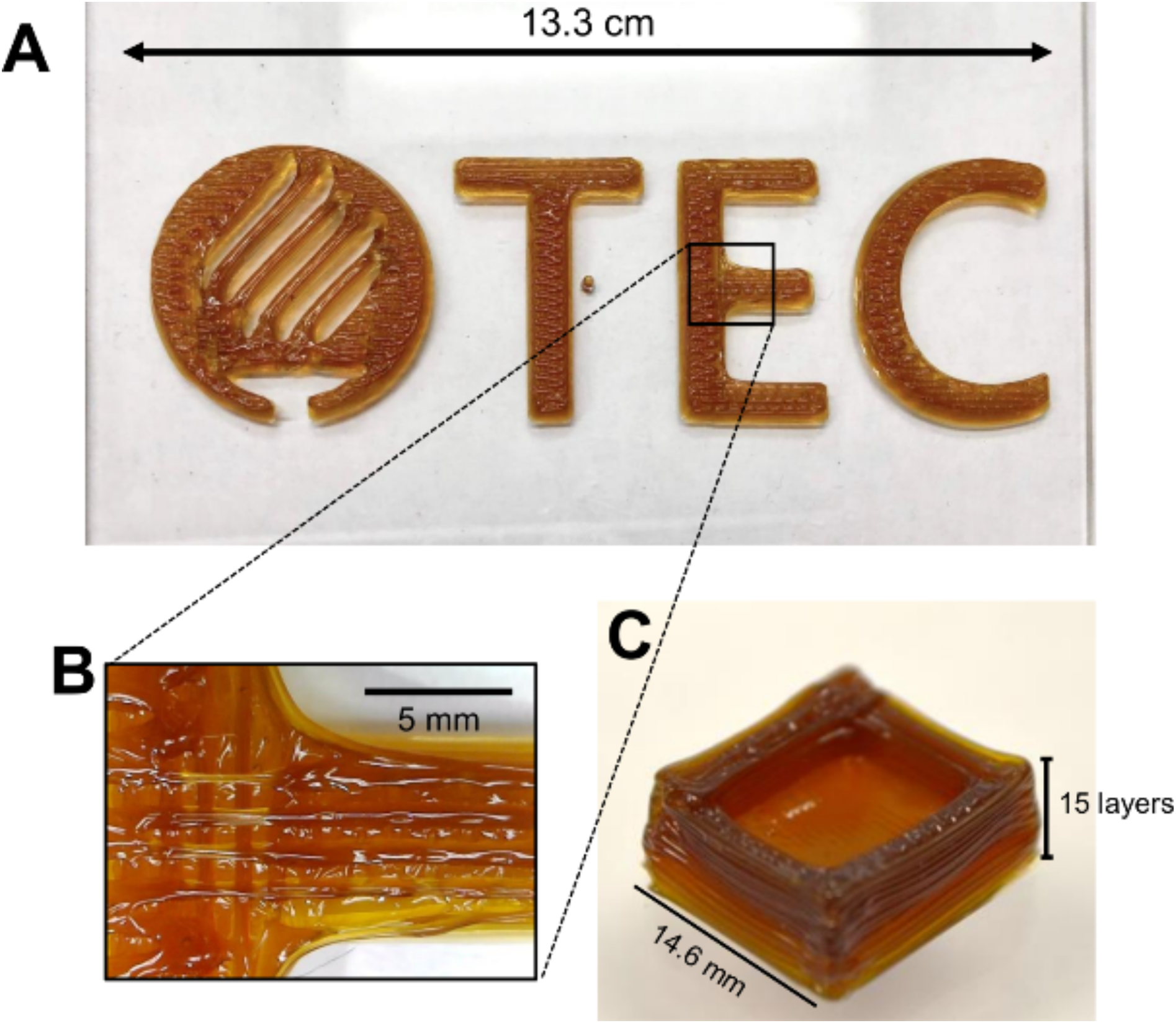
3D-printing of complex objects. A) “Tecnológico de Monterrey” logo 3D-printed with high resolution. B) Close-up of the logo showing the resolution of the printed structure. C) Self-standing hollow box printed with 15 layers. The printing parameters were 85 kPa, speed 33 mm s^−1^, and room temperature. We used a Z ink with 30 days of aging.

We further challenged the self-standing properties of this ink by printing a hollow box (Figure 5C). The large printed constructs of more than 15 layers (6 cm) did not collapse or lose their shape. Some other publications have reported printing hollow constructs of 10–15 layers; however, the studies often used sacrificial or support materials, as well as post-printing processing [45–47].

### Stability of zein-printed constructs in aqueous environments

In many biological applications, such as cell culture or drug release, printed constructs will be exposed to aqueous environments for long durations [48]. Therefore, the stability of the zein-printed constructs is a relevant property for assessment. In this section, we report the response of zein constructs to aqueous environments in terms of integrity and topographical changes. We first assessed the integrity of our printed constructs throughout typical cell culture protocols by immersing the 3D-printed zein grids in cell culture medium and incubating for 7 days at 37 °C. The average thickness of the lines of the grids and their projected surface was monitored over the 7 days (Figure 6A). The ZP grids exhibited a (non-significant) higher stability than Z grids after 7 days of culture, but both the Z and the ZP formulations were able to withstand typical cell culture conditions. For instance, the projected surface of ZP grids, as determined by image analysis of optical micrographs, was preserved by 96.2 ± 3.35%. Similarly, 92.40 ± 8.04% of the grid structure remained in the 3D-printed Z-grids after 7 days of culture. These results suggest that the extent of erosion was moderate (3.77 and 7.59% for ZP and Z respectively) after one week under culture conditions, providing evidence for stability under typical culture conditions.

**Figure 6.**
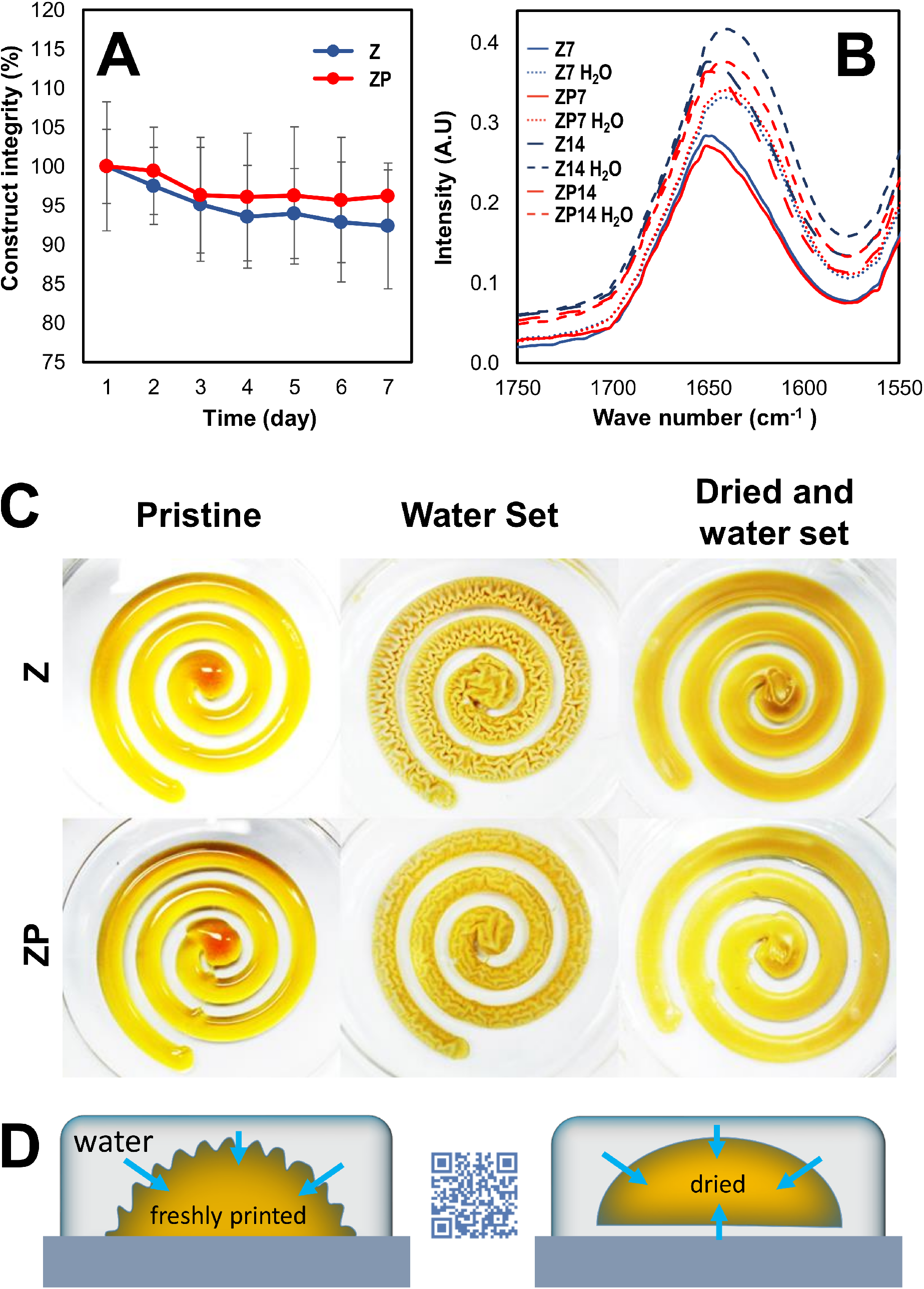
Stability of zein-printed constructs in aqueous environments. A) Z and ZP construct integrity under aqueous conditions over time. B) FTIR analysis of Z and ZP inks at two different aging times (days 7 [Z7, ZP7] and 14 [Z14, ZP14]) and after immersion in water. FTIR of samples after water immersion is indicated with the suffix H_2_O. (C) Effect of wetting freshly printed or desiccated zein-constructs on their surface topography. D) Schematic representation of the effect of swelling in zein constructs.

Zein-based materials are known to change their surface structure upon contact with aqueous solutions due to the rearrangement of their hydrophilic and hydrophobic protein domains [49]. We also investigated the effects of water exposure on the secondary structure of zein molecules in our formulations. Exposure to water can induce changes in the secondary structure of zein chains because the ethanol used as the solvent for ink formulation is displaced by water and the hydrophilic domains of zein are forced outwards [50,51]. These changes in secondary structure were evaluated by FTIR analysis (Figure 6B), which revealed a clear difference between dry and wet samples of zein-based inks. Dry samples have a symmetrical amide I band located around 1650 cm^-1^, while that band is skewed towards 1640 cm^-1^ in the wet zein.

We qualitatively investigated the changes in appearance of 3D-printed zein constructs following exposure to water. We explored two scenarios: wetting immediately after printing and wetting after 48 h of drying in a desiccator. The process of water-set causes marked topographic changes in the surface of freshly printed zein-constructs (Figure 6C). Water swelling causes very distinctive wrinkles on the surface of zein printings. Interestingly, the wrinkles are much less pronounced in ZP printings. The wrinkling effect most likely happens due to heterogeneous water swelling [49].

As shown in Figure 6D, the portion of the construct directly exposed to the aqueous environment absorbs water at a faster rate than the one that is adhered to the surface. Extrusion of zein directly into water, which equally exposes all surfaces, does not lead to wrinkling (Supplementary video 2). If 3D printed zein constructs are first desiccated for 48 hours and later immersed in water, wrinkle formation is prevented (Figure 6 C,D).

The wrinkles could have an undesired effect on the shape of printed zein constructs; however, they may be convenient for some applications. For instance, wrinkling might be exploited to increase surface/volume ratios. For example, we could envision applications where the addition of water greatly enhances the surface area available for cell attachment, particle adhesion, or mass transport enhancement at the surface. On the contrary, desiccation after printing enforces the stability and fidelity of zein printed structures and therefore their use in a wide range of biomedical applications where constructs are typically exposed to aqueous environments.

### Biomedical applications: Drug release

Zein-based materials have been used as vehicles in pharmaceutical formulations [20,52]. For instance, zein microspheres [53] and bionanocomposites [54] have been used for drug delivery. As mentioned before, zein is classified as a safe excipient by the U.S. Food and Drug Administration (FDA) [55,56].

Here, we explored the feasibility of using zein inks for 3D-printing of tablets containing a pharmaceutical compound to show that ink-based tablets may be effective alternatives for drug release applications and personalized therapies. We used the NA antibiotic as a pharmaceutical model in our controlled release experiments from zein-printed tablets, as NA is a first-generation quinolone-based compound that is able to mitigate narrow spectrum implant-related infections [56–58]. Figure 7A shows results of a 48 h drug-release experiment from zein-based printed tablets containing NA. A burst effect was observed for Z and ZP during the first 8 h, with a release of 29.83 ± 0.87% and 39.90 ± 2.04% of the drug, respectively. The incorporation of PEG400 enhanced the burst effect by 10%, arguably because zein chains are less entangled and allow faster diffusion of the NA through the polymeric matrix. The hydrophilicity of PEG400 may also play an important role in the diffusion of NA, as shown elsewhere for different pharmaceutics and drug delivery materials [60,61]. At 48 h, a significantly higher NA load was released from the ZP tablet formulations (93.84 ± 2.98%) than from the Z formulations (65.49 ± 1.73%).

**Figure 7.**
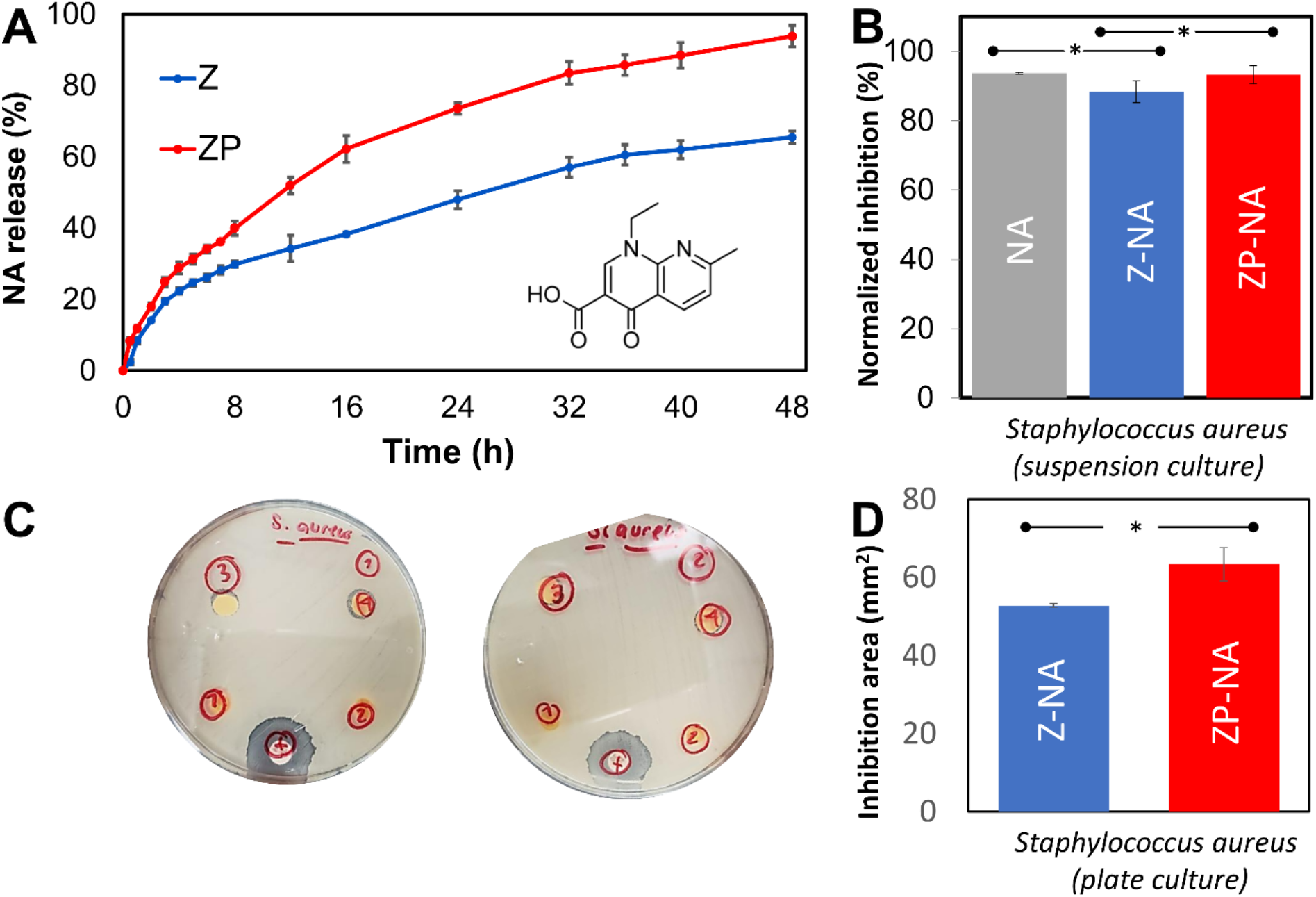
Drug release. A) Cumulative release of NA from Z (blue line) and ZP (red line) for 48 h (chemical structure of NA is shown on the bottom right). (B) Bacterial inhibition by 3D-printed tablets of Z-NA and ZP-NA against suspension cultures of *Staphylococcus aureus*. (C-D) Bacterial inhibition by 3D-printed tablets of Z-NA and ZP-NA against suspension cultures of *Staphylococcus aureus* in plate cultures. (C) Images of two independent replicas are presented. The imbibition halos induced by a disc of NA (as a positive control) (+), 3D-printed tablets of Z-NA (1,2), and ZP-NA (3,4) are shown, and (D) their area was quantified by image analysis.

We also determined the antibacterial activity of 3D-printed zein tablets formulated with NA against suspension cultures of *Staphylococcus aureus* (Figure 7B). Four different tablet compositions (pristine Z, pristine ZP, Z with added NA [Z-NA], and ZP with added NA [ZP-NA]) were evaluated. Control experiments with direct NA application (without zein) showed that the NA drastically inhibits the growth of *S. aureus* at the concentrations tested. We observed a decrease of 93.60% in the absorbance of *S. aureus* suspension cultures, in agreement with previous reports [62,63].

We also evaluated the antibacterial effect of 3D-printed Z and ZP tablets containing a controlled-release dose of 3 mg NA per g of zein. We observed a decrease of 88.27% in absorbance of the bacterial culture after 48 h in experiments where Z-NA 3D-printed tablets were added to *S. aureus* suspension cultures (Figure 7B) and plate cultures (Figure 7C-D). The antibacterial activity was even higher for the ZP-NA tablets, which caused a 93.20% decrease in the absorbance of *S. aureus* suspension cultures. ZP-NA 3D-printed tablets also induced significantly larger inhibition halos than Z-NA tablets in solid agar cultures 48 hours after plating highly concentrated *S. aureus* suspensions (10^7^ cells mL^−1^).

### Cellular scaffolding

We also explored the feasibility of using 3D-printed zein scaffolds to support cell cultures. We evaluated cell attachment and morphology using optical and fluorescence microscopy (Figure 8A,B) and cell viability (using the PrestoBlue^®^ assay) over time in C2C12 cells cultured on zein surfaces (Figure 8C). We observed that cell attachment and spreading was not favored on untreated 3D-printed Z and ZP surfaces.

**Figure 8.**
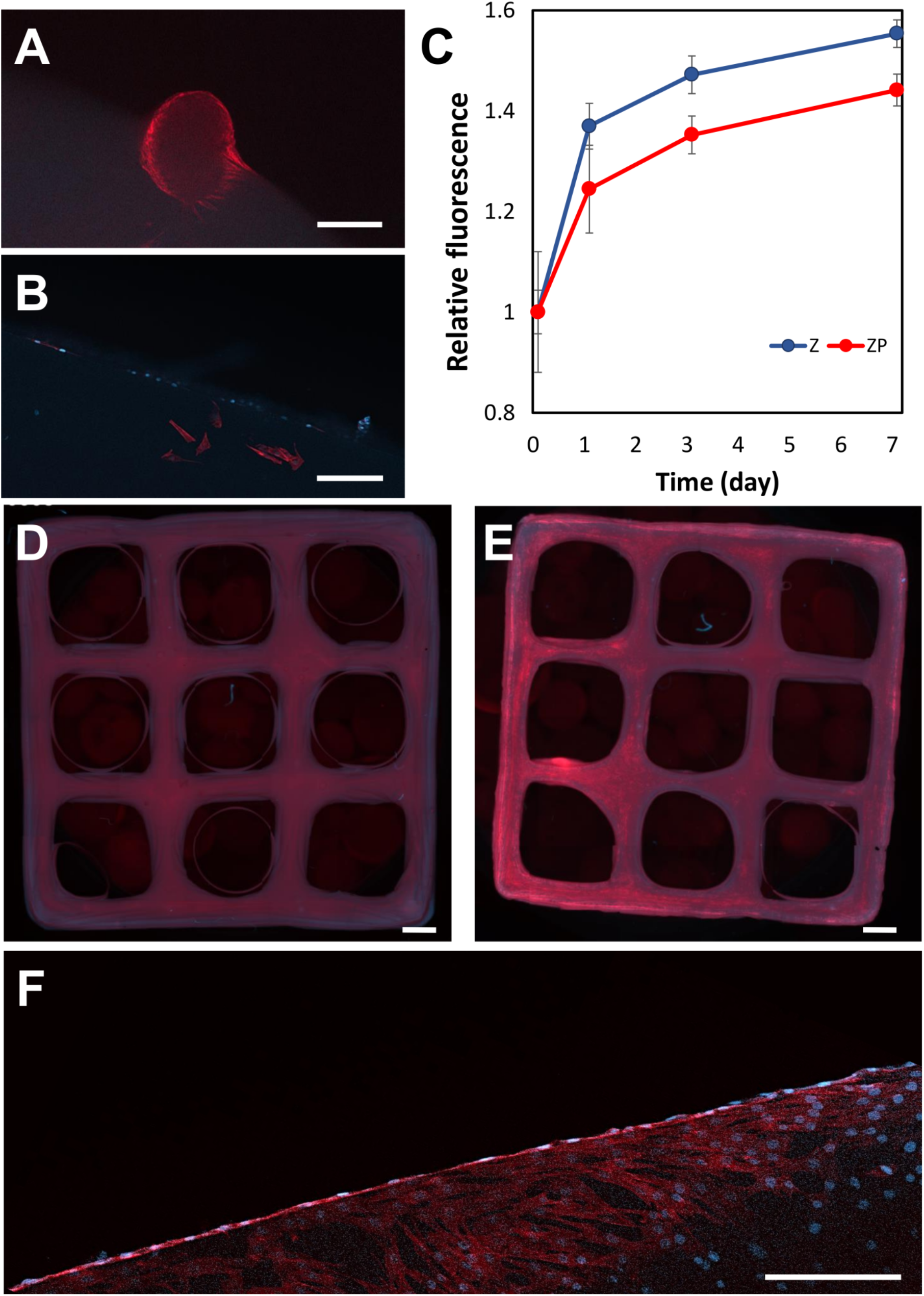
Zein-based 3D printed constructs as cellular scaffolds. A) Actin-stained C2C12 agglomerate formed on the surface of a grid printed with Z at day 3 of culture; scale bar: 200 μm. B) Actin/DAPI staining of C2C12 cells at day 7 of culture on a ZP grid; scale bar: 200 μm. C) Normalized C2C12 myoblast Prestoblue proliferation assay on the 3D-printed grids of Z (blue) and ZP (red) over 7 days of culture. D) Actin/DAPI staining of C2C12 cells at day 7 of culture on a ZP grid; scale bar: 2000 μm. E) Actin/DAPI staining of C2C12 cells at day 7 of culture on ZP grids covered with fibronectin; scale bar: 2000 μm. F) Close up of cells spreading at day 7 of culture on ZP grids covered with fibronectin, scale bar: 200 μm.

For instance, Figure 8A shows only limited cell attachment on Z-printed constructs, as revealed by the tendency to form agglomerates of cells (colonies) instead of a homogenous monolayer over the surface. Note, however, that the tissue spheroids anchored to Z constructs could be useful in organ-on-chip scenarios, such as anti-cancer drug screening using anchored cancer spheroids. Figure 8B also shows very low cell attachment to ZP-printed constructs. The metabolic activity of C2C12 cells was higher when cultured on 3D-printed Z grids than on ZP grids (Figure 8A-C). Taken together with the metabolic activity data, the microscopy examination indicates that zein-based materials are cytocompatible but provide limited anchorage capacity (zein does not exhibit RGD sequences) for cell attachment.

We conducted an additional set of cell culture experiments using 3D-printed zein-based grids coated with fibronectin, a widely-used cell-adhesion protein [64]. We then compared actin/DAPI fluorescent stainings on grids of ZP (Figure 8D) and ZP with fibronectin (Figure 8E) after 7 days of C2C12 cell culture. The close-up micrographs of the ZP with fibronectin grids (Figure 8F) showed that the fibronectin coating significantly enhanced cell attachment and spreading. Overall, these results provide evidence of the potential for use of 3D-printed zein-based materials as cell culture substrates.

## Conclusions

The current portfolio of 3D-printable proteins is limited. Here, we introduced and characterized 3D-printing inks based on zein, which is the main constitutive protein in maize seeds and a highly available and low-cost material. These inks are easily prepared with standard laboratory tools and their rheology, and thus printability, can be tuned by altering the aging time and/or printing temperature or by the addition of plasticizers such as PEG. Printability was analyzed for three different printing pressures and three different printhead velocities. These parameters can be adjusted to achieve a desired resolution depending on the different rheological properties of the inks. Remarkably, unlike other protein-based inks, our zein inks generated 3D-printed structures with no requirement for either a pre-chemical functionalization or a post-stabilizing process. These inks are therefore highly versatile for use in 3D printing applications.

Knowing the range of capacities of a particular bioprinter, one can choose an adequate formulation for a particular profile of pressures and speeds. Even for a fixed composition, one can tune the rheology of a zein-based ink by manipulating the maturation times and PEG concentrations. Proof-of-concept studies confirmed that zein-bioprinted constructs can used for drug-release applications. We also explored the use of zein inks in the fabrication of 3D printed cell scaffolds. Zein-based grids were printed, coated with fibronectin, and seeded with C2C12 myoblast cells. We observed successful cell attachment, spreading, and viability over a period of 7 days. Zein is recognized as safe by the FDA; therefore, 3D printing with zein based inks will open the door to versatile applications in diverse fields and especially biomedical applications.

## Acknowledgments

JANT, PATC. And JMOC gratefully acknowledge the finantial support received from CONACyT (Consejo Nacional de Ciencia y Tecnología, México) in the form of Graduate Program Scolarships. ASM, GTdS, and MMA acknowledge the institutional funding received from Tecnológico de Monterrey (Grant 002EICIS01). MMA, GTdS acknowledge funding provided by CONACyT (Consejo Nacional de Ciencia y Tecnología, México) through grants SNI 26048, SNI 256730, and SNI 1056909, respectively. AT would like to acknowledge the financial support of National Institutes of Health (GM126831, AR073822)

