## Supplementary File for "Biofabrication using maize protein: 3D printing using zein formulations"

### Supplementary information

#### FTIR analysis

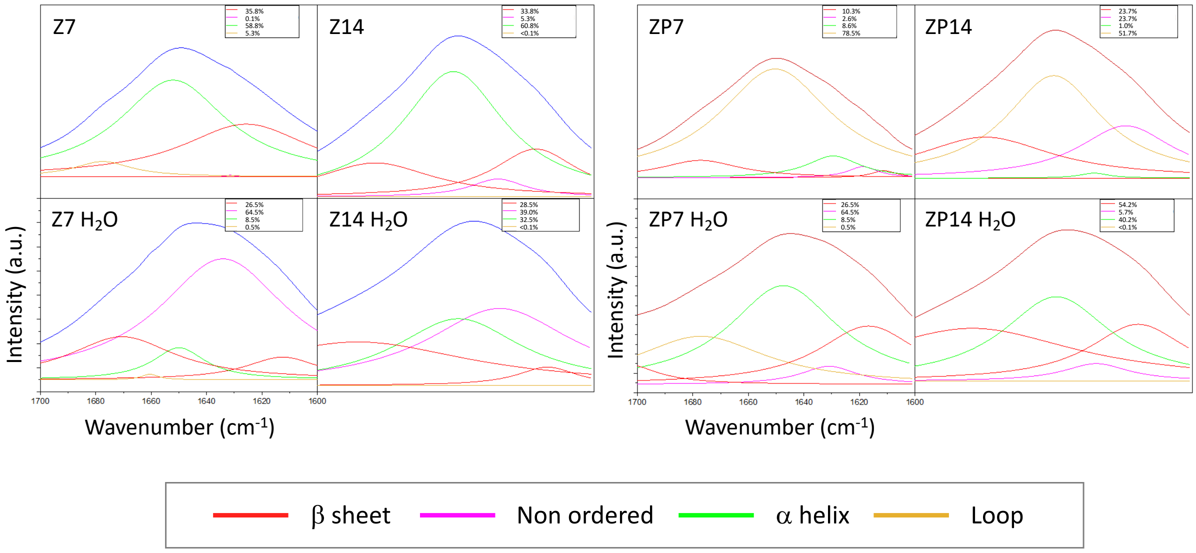


**Figure S1. Deconvolution of the amide I peak** of Z and ZP inks at different aging time points (day 7 and day 14) and before and after immersion in water ($H_2O$).

**Table 1. Quantitative data from the deconvolution of FTIR in Figure SI 1**

| Secondary structure | Zein | | | | Zein + PEG | | | |
|---|---|---|---|---|---|---|---|---|
| | Dry | | Wet | | Dry | | Wet | |
| | Day 7 | Day 14 | Day 7 | Day 14 | Day 7 | Day 14 | Day 7 | Day 14 |
| β-sheet | 35.84% | 33.84% | 26.48% | 28.51% | 10.31% | 23.65% | 24.99% | 54.15% |
| Non-ordered | 0.09% | 5.31% | 64.46% | 39.04% | 2.60% | 23.68% | 5.17% | 5.66% |
| α-helix | 58.79% | 60.85% | 8.53% | 32.45% | 8.64% | 0.95% | 46.61% | 40.19% |
| Loop | 5.28% | <0.01% | 0.53% | <0.01% | 78.45% | 51.72% | 23.23% | <0.01% |

**Internal microstructure of extruded zein inks**

We conducted image analysis to characterize the microstructural changes in zein formulations through time (Figure 4A). With that aim, we extruded lines of zein-based inks, broke them by fast freezing in liquid nitrogen, and determined the size distribution of pores within cross-sections. In general, the number of pores and pore area decreased with increasing aging time. As expected, the plasticity provided by PEG400 resulted in low-porosity values for ZP formulations. Consistently, the cross-sections had smoother surfaces and smaller average pore sizes for ZP than for Z samples. Figure 4B shows that 73.5% of the pores in 7-day aged Z ink had pore sizes between 0.2 and 1.26 $\mu m^2$ (the first bin), whereas 81.8% of in pores in 14-day aged Z ink had measured pore sizes between 0.2 and 1.26 $\mu m^2$, while 92.24% of the pores were in the smallest bin in 21-day aged Z. Conversely, for ZP, the percentages of pores smaller than 0.83 $\mu m^2$ on days 7, 14, and 21 were 41.73%, 76.85%, and 86.73%, respectively.

Note also that the pore area deviation decreased as maturation time progressed, suggesting that the pore sizes become more homogeneous over time.

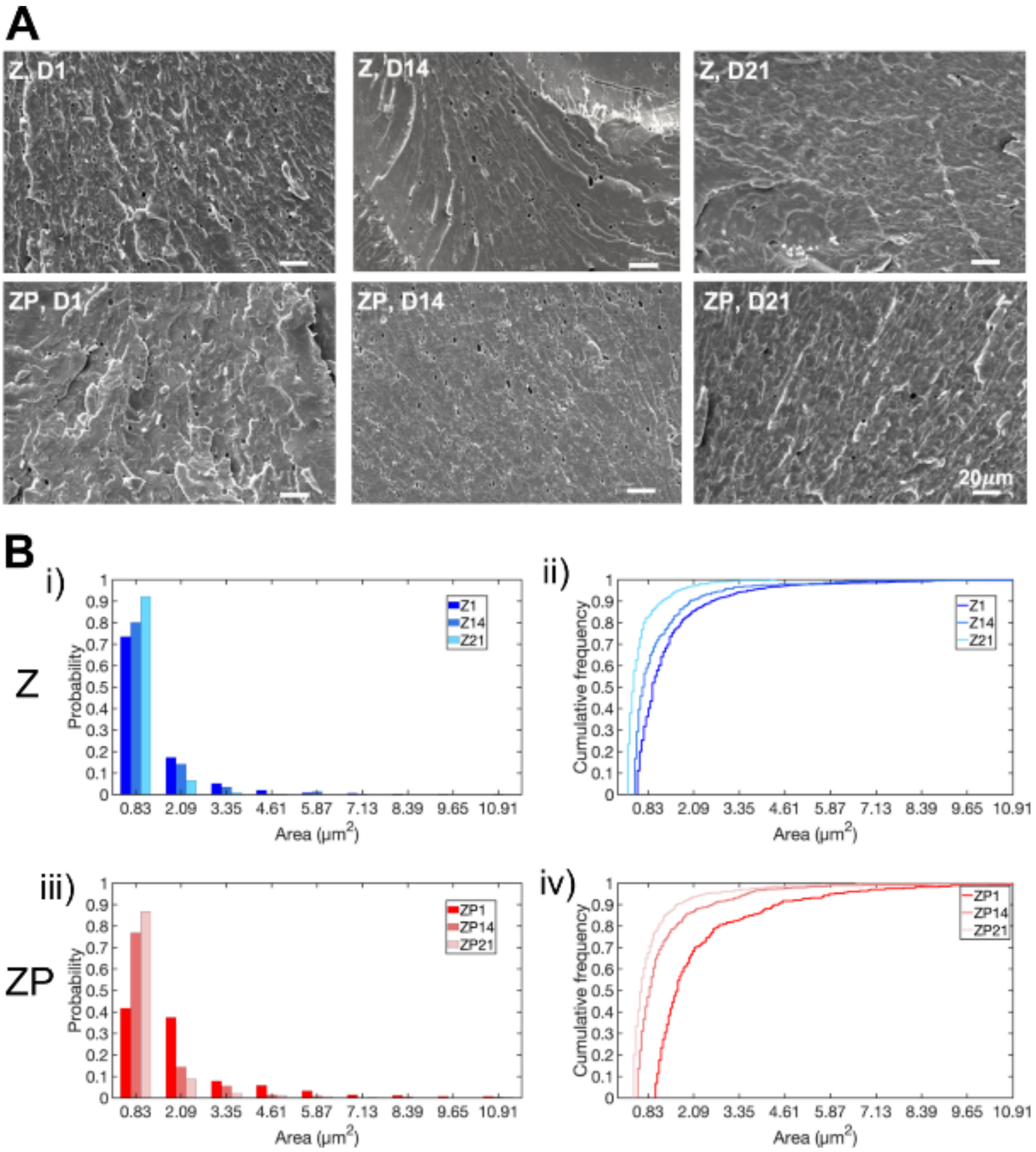


**Figure SI 2. SEM pore analysis of the cross-sections of printed constructs.** A) Micrographs from Z and ZP inks at days 1, 14, and 21. Scale bar: 20 μm. B) Histogram distribution of pore sizes in Z and ZP across time.

Supplementary video 1: Zein 6 layer construct (with infill)

https://youtu.be/RW6c7JsWMZQ

Supplementary video 2: Zein 10 layer construct (open triangle)
https://youtu.be/SVJrhlVFkb0

Supplementary video 3: Water setting

https://youtu.be/shUPkXQj5FA
